# Neuronal patterning of the tubular collar cord is highly conserved among enteropneusts but dissimilar to the chordate neural tube

**DOI:** 10.1101/118612

**Authors:** Sabrina Kaul-Strehlow, Makoto Urata, Daniela Praher, Andreas Wanninger

**Affiliations:** Department for Integrative Zoology, University of Vienna, Althanstr. 14, 1090 Vienna, Austria; Research Center for Marine Biology, Tohoku University, Asamushi, Aomori, Aomori 0393501, Japan; Noto Marine Laboratory, Division of Marine Environmental Studies, Institute of Nature and Environmental Technology, Kanazawa University, Ogi, Noto-cho, Ishikawa 927-0553, Japan

## Abstract

The dorsal neural tube of chordates and the ventral nerve cord of annelids exhibit a similar molecular mediolateral architecture. Accordingly, the presence of such a complex nervous system (CNS) has been proposed for their last common ancestor. Members of Enteropneusta, a group of non-chordate deuterostomes, possess a less complex CNS including a hollow neural tube, whereby homology to its chordate counterpart remains elusive. Since the majority of data on enteropneusts stem from *Saccoglossus kowalevskii,* a derived direct-developer, we investigated expression of key neuronal patterning genes in the indirect-developer *Balanoglossus misakiensis.*

The collar cord of *B. misakiensis* shows anterior *Six3/6* and posterior *Otx* + *engrailed* expression, in a region corresponding to the chordate brain. Neuronal *Nk2.1/Nk2.2* expression is absent. Interestingly, we found median *Dlx* and lateral *Pax6* expression domains, i.e., a condition that is reversed compared to chordates.

Comparative analyses reveal that CNS patterning is highly conserved among enteropneusts. *BmiDlx* and *BmiPax6* have no corresponding expression domains in the chordate brain, which may be indicative of independent acquisition of a tubular CNS in Enteropneusta and Chordata. Moreover, mediolateral architecture varies considerably among chordates and enteropneusts, questioning the presence of a vertebrate-like patterned nervous system in the last common deuterostome ancestor.

## Introduction

The evolution of the bilaterian central nervous system (CNS) has been hotly debated for decades [1–6]. In this debate, enteropneust hemichordates (or acorn worms) have occupied a pivotal role. Enteropneusts are a group of hemichordate deuterostomes distantly related to vertebrates, which have retained a number of putative ancestral bilaterian features such as a biphasic life style and a bilateral symmetric body. Enteropneusts share some characteristics with chordates, such as gill slits and, at least partly, a tubular nervous system. For these reasons, enteropneusts are ideal candidates to unravel the evolution of the nervous system of Deuterostomia. The majority of enteropneust species belong to one of the three main families Harrimaniidae (e.g. *Saccoglossus kowalevskii)*, Speneglidae (e.g. *Schizocardium californicum)* and Ptychoderidae (e.g. *Balanoglossus* spp., *Ptychodera flava)* [7]. Harrimaniid species develop directly into the juvenile worm, whereas spengelid and ptychoderid enteropneusts develop indirectly via a specific larval type, the tornaria. Morphologically, the nervous system of enteropneusts is described as a basiepidermal plexus with additional condensed regions. These comprise the proboscis stem and nerve ring, a dorsal nerve cord along the collar and trunk region, and a ventral nerve cord in the trunk connected to the dorsal nerve cord by a prebranchial nerve ring (Fig.1A’) [8, 9]. The dorsal nerve cord within the collar region, the ‘collar cord’, is a subepidermal tubular nerve cord that is often thought to be reminiscent of the chordate neural tube and, like the latter, forms by neurulation [10, 11]. The collar cord is subdivided into a dorsal sheath of different neuronal cell types surrounding a central neural canal and a ventral neuropil [11, 12]. Although these morphological features would support homology of the chordate neural tube and the collar cord of enteropneusts, it remains unclear to which part of the chordate neural tube the collar cord might correspond to. Moreover, the results from gene expression analyses are somewhat contradictory. The CNS of many bilaterians is patterned similarly from anterior to posterior by a number of specific transcription factors (see [3] for review). For instance, genes such as *Six3/6, Otx* and *engrailed* regionalize different parts of the brain in bilaterians, while *Hox* genes pattern the postcerebral region (spinal cord or ventral nerve cord, respectively). Anteroposterior patterning of these transcription factors has been studied in the enteropneust *S. kowalevskii* and is similar to that in chordates [3, 13, 14], yet the expression domains in enteropneusts are circumferential in the entire ectoderm and not restricted to the CNS-forming domains (“neuroectoderm”) as in chordates [13, 14]. Like in *S. kowalevskii,* a recent study described similar expression domains of those transcription factors in the spengelid *S. califoricum* [15]. To complicate things further, Miyamoto and Wada [16] showed that genes specifying the chordate neural plate border (e.g., *SoxE,* and *Bmp2/4*) have conserved expression domains in the collar cord of the enteropneust *Balanoglossus simodensis* [16]. Concluding so far, no unequivocal homology statement can be made at present concerning the collar cord and the chordate neural tube.

**Fig. 1.**
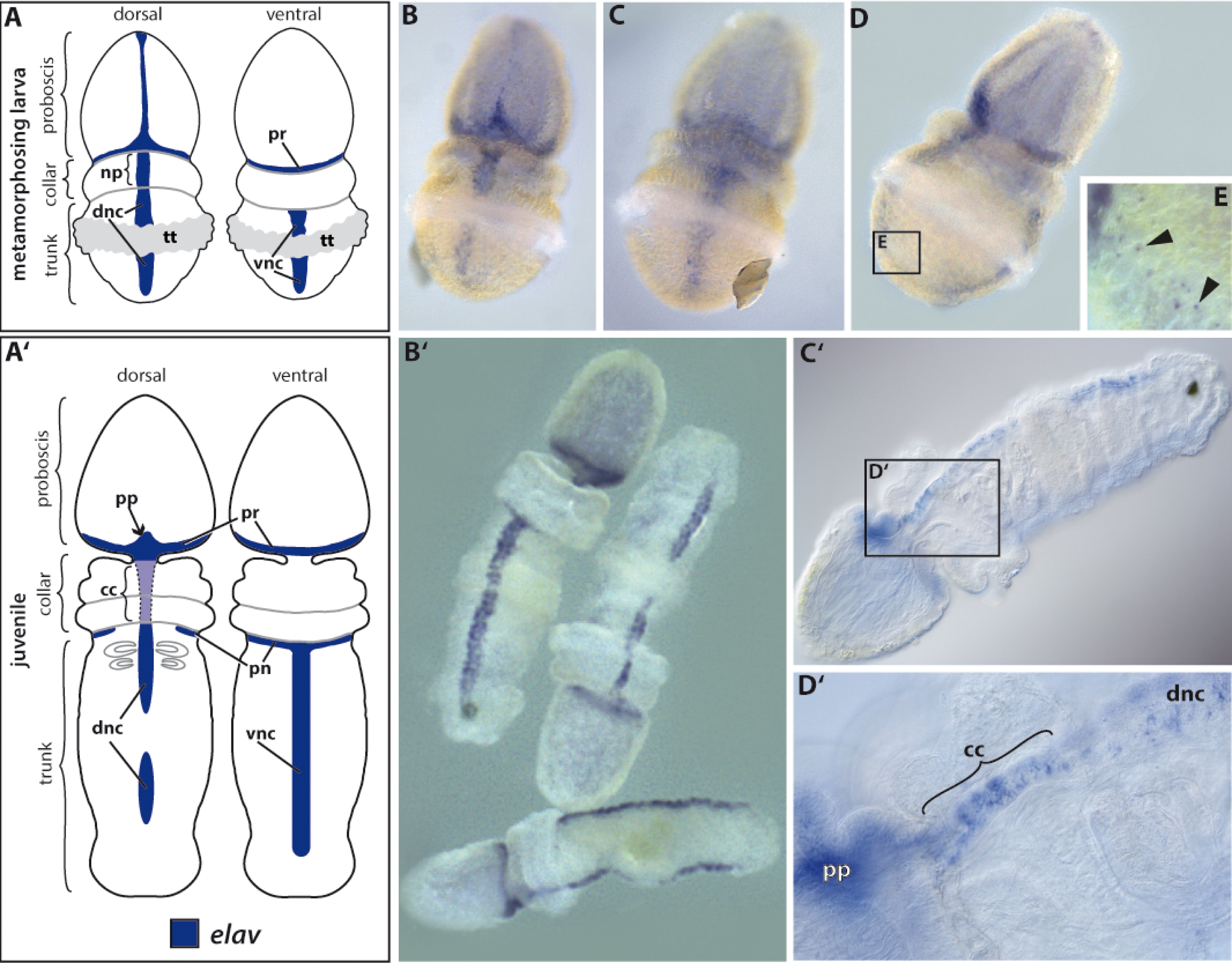
Establishment of the adult nervous system. Gene expression of *BmiElav* in the metamorphosing larva and juvenile of *B*. *misakiensis*. **A-E** Metamorphosing larva. **A’-D’** Juvenile. **A** Schematic illustration of *BmiElav* expression. *BmiElav* is expressed in the proboscis nerve ring, the developing dorsal nerve cord including the neural plate in the collar (**B, D**) and in the ventral nerve cord (**C, D**). Note that the expression is interrupted at the level of the telotroch. **E** Detail of the lateral trunk showing scattered neurons (arrowheads). **A’** Schematic illustration of *BmiElav* expression in juveniles. Note that the collar cord is neurulated. **B’** Surface view from ventral, dorsal and lateral right (from top to bottom) showing strong expression in the proboscis nerve ring, proboscis plexus, and in the dorsal as well as ventral nerve cord. *BmiElav* expression is discontinuous in the middle of the dorsal nerve cord in the trunk region. **C’** Micrograph of cleared juvenile. **D’** Detail showing *Elav+* cells in the subepidermal collar cord. cc = collar cord, dnc = dorsal nerve cord, np = neural plate, pn = peribranchial nerve ring, pr = proboscis nerve ring, pp = proboscis plexus, tt = telotroch, vnc = ventral nerve cord. B dorsal view. C ventral view. D view from lateral right.

Most of the molecular data available for enteropneusts have been obtained from *S. kowalevskii,* a harrimaniid with derived direct development. This data has been supplemented recently by a bodypatterning study of the spengelid *S. californicum* [15], yet comparable data from a ptychoderid species are still missing. However, a reliable ground pattern of neuronal patterning for Enteropneusta can only be reconstructed, if data from members of as many different enteropneust families are compared. Thus, in order to contribute new insights into the evolution of tubular nervous systems in Deuterostomia a comparable study on neuronal patterning of the collar cord in a ptychoderid enteropneust is of prime importance.

Here, we studied the expression domains of neuronal patterning genes in the indirectly developing ptychoderid *Balanoglossus misakiensis*, in order to provide the missing data. We focused on the expression patterns in the developing collar cord of anteroposterior (*Six3/6, Otx,* and *engrailed*) as well as so-called mediolateral patterning genes (*Pax6, Dlx, Nk2.1, Nk2.2*). The latter have been reported to form abutting domains of *Nk* and *Pax* genes in the annelid ventral nerve cord and in the vertebrate dorsal neural tube [2, 17]. In each of these progenitor domains specific neuronal cell types are formed. For instance, serotonin-positive (+) neurons are exclusively restricted to the median *Nk2.1* domain in the brain and to the median *Nk2.2* in the trunk nerve cord [2]. *Pax6* forms two bilaterally symmetric, intermediate progenitor domains and *Dlx* two lateral domains. Given the complexity of the corresponding spatial organization of the annelid nerve cord and the vertebrate neural tube, a similarly patterned nervous system has been proposed in the Urbilaterian. Herein, we assess the presence of putative mediolateral patterning in the collar cord of *B. misakiensis*. This study comprises the first gene expression data for this particular species. Our data allow for a comparison with other enteropneusts and lead to a reliable ground pattern reconstruction for Enteropneusta. Eventually, this will help comparing the collar cord to the chordate neural tube and contribute to our understanding of nervous system evolution in Deuterostomia.

## Results

### Neuronal differentiation of the adult nervous system

The central nervous system of *B. misakiensis* including the collar cord become morphologically distinct in early settled juveniles, indicating that neurogenic patterning of the collar cord starts in metamorphosing larvae [18]. In contrast, the larval nervous system (apical organ and neurite bundles of the ciliary bands) are independent of the adult nervous system and degrade during metamorphosis and settlement [7, 16, 19]. Therefore, we focused on the expression patterns in metamorphosing larvae and early juveniles.

In order to obtain an overview of the developing adult nervous system of *Balanoglossus misakiensis,* we first examined the expression of *Elav,* an RNA-binding protein that marks differentiating neurons [20–22]. *BmiElav* is expressed in the epidermis of the metamorphosing larva (Agassiz stage) of *B. misakiensis* as a stripe along the entire dorsal midline (except at the level of the telotroch) and extends circumferentially to the posterior base of the proboscis (Fig. 1A, B, D). In addition, *BmiElav* expression runs along the ventral midline of the trunk region with a gap in the region of the telotroch (Fig. 1A, C, D). *BmiElav* thus includes the region of the future dorsal and ventral nerve cords. Higher magnification of the perianal field reveals additional scattered *BmiElav*+ cells laterally outside the nerve cords (Fig. 1E).

In juvenile *B. misakiensis, Elav+* cells are abundant in all condensed parts of the nervous system [18], including the proboscis plexus at the base of the proboscis region and the proboscis nerve ring (Fig. 1 A’, B’). At the level of the collar region, *BmiElav+* cells locate to the subepidermal collar cord (Fig. 1D’). *BmiElav+* cells are also present in the prebranchial nerve ring, as well as in the dorsal and ventral nerve cords in the trunk region (Fig. 1B’, C’). The *BmiElav* signal is interrupted in the dorsal nerve cord at the former position of the telotroch.

### Gene expression of anteroposterior patterning genes

We studied the expression of selected axial patterning genes in order to determine the region to which the enteropneust collar cord might correspond to in the vertebrate neural tube.

The transcription factor *BmiSix3/6* is strongly expressed throughout the entire ectoderm of the proboscis region and extends into the anterior rim of the collar ectoderm in metamorphosing larvae (Fig. 2A, B) and juvenile worms (Fig. 2C, D).

**Fig. 2.**
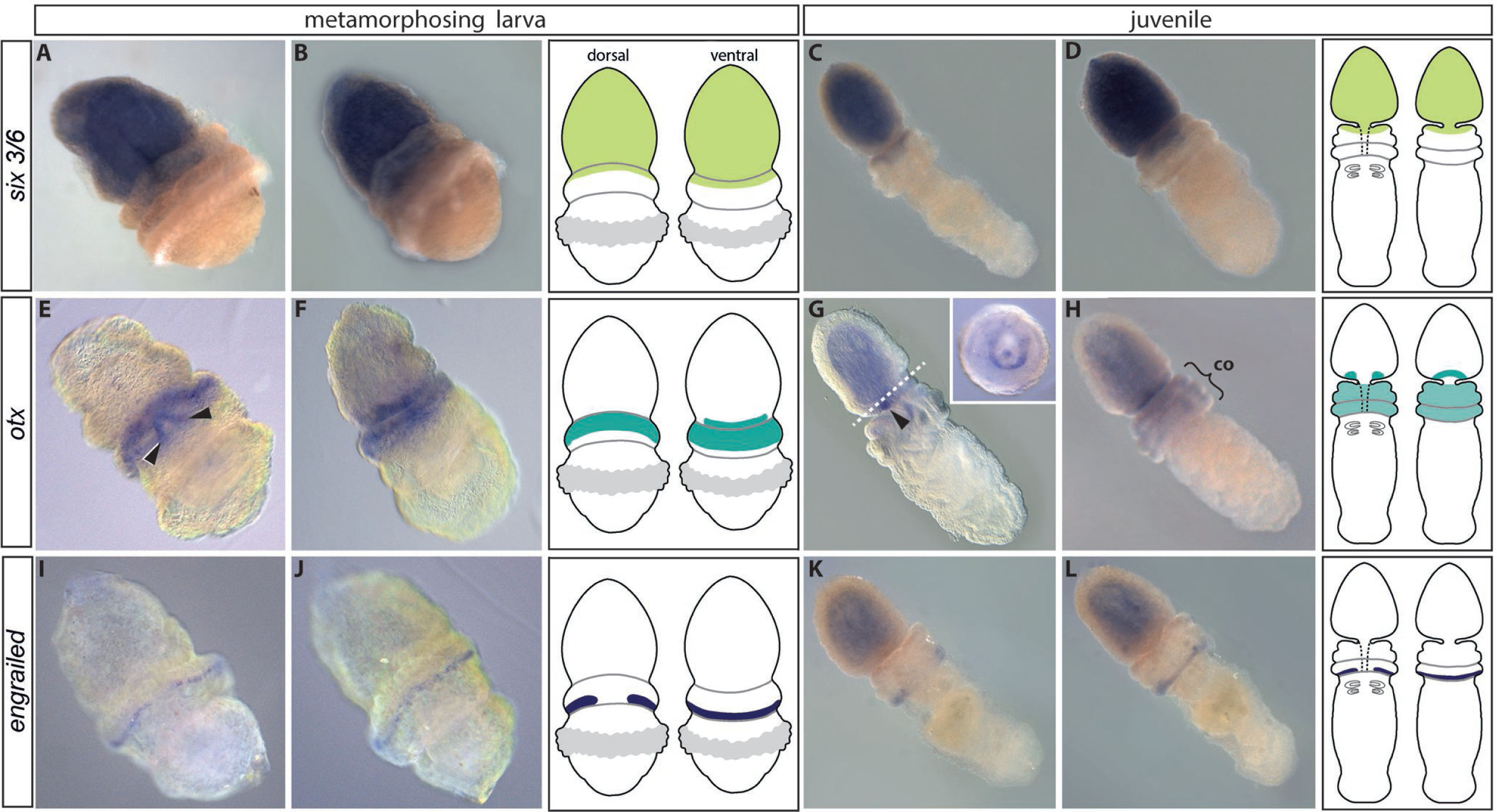
Anteroposterior patterning genes allocate the collar cord of *B. misakiensis* to the chordate brain region. Anterior is to the top left. **A-D** *BmiSix3/6* is expressed throughout the ectoderm of the proboscis region and the anterior collar. **E-H** *BmiOtx* is expressed circumferentially in the posterior proboscis ectoderm and in the ectoderm of the collar region. **E** Dorsal view showing an additional domain in the pharyngeal endoderm (arrowheads). **G** *BmiOtx* is strongly expressed in the preoral ciliary organ (arrowhead). Section plane of inset indicated by dashed line. Inset shows *Otx* expression in the ciliary organ of the proboscis in a cross section of the posterior proboscis. **I-L** *BmiEn* is expressed in a narrow ring in the ectoderm of the posterior end of the collar region with an interruption on the dorsal side. co = collar. A, C, E, G, I, K: dorsal views. B, D, F, H, J, L: ventral views.

*BmiOtx* is expressed in the metamorphosing larva in the ventral area of the proboscis nerve ring (Fig. 2F) and in a distinct annular domain, encircling the anterior and middle collar region (Fig. 2E, F, Fig. S1B). The additional domain in the anterior pharyngeal region (Fig. 2E arrowheads) is a non-neural endodermal domain that will not be further discussed. In the juvenile enteropneust *BmiOtx* is expressed in the ventral and ventrolateral area of the proboscis nerve ring (Fig. 2G, H). The expression forms a U-shaped domain at the position where the sensory ‘pre-oral ciliary organ’ develops (Fig. 2G inset). *BmiOtx* is also weakly expressed throughout the ectoderm of the collar region (Fig. 2H). The signal within the proboscis coelom is due to probe trapping and interpreted as unspecific background (Fig. S1). *BmiEn* is expressed in a circumferential ring at the very posterior margin of the collar region in metamorphosing larvae (Fig. 2I, J and Fig. S1C). The signal is ectodermal and interrupted at the level of the dorsal midline. The juvenile enteropneust shows a similar expression pattern at the posterior margin of the collar region (Fig. 2K, L). The ring of *BmiEn* expression shows a gap on the dorsal side, as in the metamorphosing larva.

In summary, the collar cord, that is part of the enteropneust collar region (mesosome), abuts anteriorly the expression domain of *BmiSix3/6*, lies within the *BmiOtx-expression* region, and is posteriorly delimitated by a line of *BmiEn* expression.

### Gene expression of mediolateral patterning genes

In metamorphosing larvae, *BmiPax6* is strongly expressed in the proboscis nerve ring at the base of the proboscis and in an additional circular pattern in the ectoderm of the posterior collar region (Fig. 3A, B). Between both circumferential domains, *BmiPax6* is also expressed in two parallel, longitudinal domains of the collar (Fig. 3A, dashed area). This area of the neural plate will later neurulate to form the subepidermal collar cord [11]. In juveniles, *BmiPax6* still shows a strong signal in the proboscis nerve ring. The circular domain in the posterior collar region becomes fainter in early juveniles (Fig. 3C inset) and is lost in older juveniles (Fig. 3C, D). No collar cord *BmiPax6* expression domains are present in juveniles. Expression of *BmiDlx* is present in the proboscis nerve ring and along the dorsal nerve cord with an interruption at the level of the telotroch in the metamorphosing larva (Fig. 3E, F). In juveniles of *B. misakiensis Dlx* expression shows a faint signal in the ventral and ventrolateral portion of the proboscis nerve ring and in the dorsal nerve cord including the collar cord (Fig. 3G, H). Our data show that *BmiDlx* is expressed in the collar cord and in the dorsal nerve cord and forms a single median domain.

**Fig. 3.**
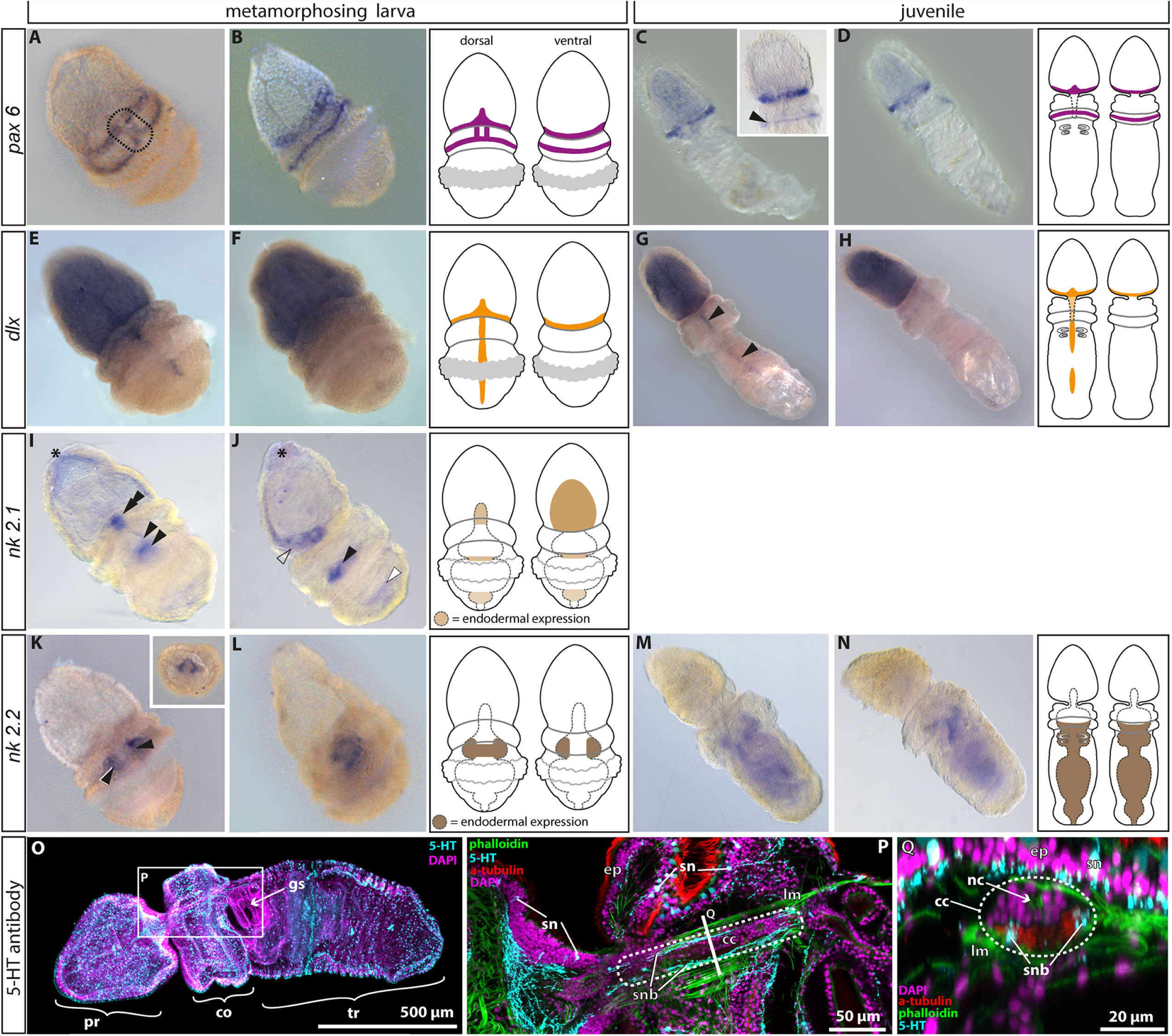
Expression domains of mediolateral patterning genes and serotonin-LIR in *B misakiensis.* **A, B** *BmiPax6* is expressed in the proboscis nerve ring and in a second circumferential domain in the collar ectoderm. Additionally, *BmiPax6* forms paired longitudinal domains in the neural plate (dashed area) of the developing larva. In juveniles, the expression in the collar ectoderm fades (inset) and only the proboscis nerve ring shows strong signal of *BmiPax6* (**C, D**). **E-H** *BmiDlx* is expressed as a median stripe in the collar and dorsal cord (arrowheads) with an interruption at the level of the telotroch (E). The strong staining in the protocoel is unspecific (see also Fig. S1). **I, J** Expression of *BmiNk2.l* in the metamorphosing larva. *BmiNk2.l* is strongly expressed in the stomochord (double arrowhead), in the ventrolateral ectoderm at the base of the proboscis (open arrowhead), in the posterior pharynx (black arrowhead), and weakly in the hindgut (white arrowhead). The degrading apical organ shows a faint signal (asterisk). **K-N** Expression of *BmiNk2.2*. **K** Surface view from ventral showing bilateral domains in the mid-pharynx region. **L** Lateral view from left. Inset shows a cross section of the collar region with the ventrolateral domain of *BmiNk2.2* in the pharyngeal endoderm. In juveniles the expression domain of *Nk2.2* is extended throughout the entire endoderm (**M, N**). Note that there is no ectodermal or neuronal expression domain of *BmiNk2.2*. **O-Q** Serotonin-LIR in the juvenile. **O** Overview. **P** Detail of the collar region as indicated in G. Partial Z-projection focussing on the collar cord (dashed area). **Q** Virtual cross section of the collar cord as indicated in H. Note that 5-HT+ somata are absent from the collar cord (dashed area). Only two ventrolateral 5-HT+ neurite bundles pass the neuropil. cc = collar cord, co = collar, ep = epidermis, g s= sill slit, lm = longitudinal muscles, nc = neural canal, pr = proboscis, sn = serotonin-LIR neuron, snb = serotonin-LIR neurite bundle, tr = trunk. A, C, E, G, I, K, M: dorsal views, B, J, L, N: lateral views left, D, F, H: ventral views.

*BmiNkx2.1* has four distinct expression domains in the metamorphosing larva (Fig. 3I, J). At this stage, the apical organ is degrading [18] and *BmiNkx2.1* is weakly expressed in this ectodermal region (Fig. 3I, J asterisk). *BmiNkx2.1* shows a strong expression domain in the ventral ectoderm at the base of the proboscis region (Fig.3J unfilled arrowhead). Further strong domains are within the developing endodermal stomochord (Fig. 3I, double arrowhead) and medially in the posterior pharyngeal endoderm (Fig. 3I, J black arrowhead). A fifth, although weak signal, is present in the hindgut (Fig. 3J, white arrowhead).

The transcription factor *BmiNkx2.2* is strongly expressed in the lateral and dorsal portions of the anterior pharyngeal endoderm in the metamorphosing larva (Fig. 3K, inset, L). In the juvenile worm the *BmiNkx2.2* domain has extended posteriorly and is present throughout the endoderm, but absent from the hindgut (Fig. 3M, N). Thus, there is no expression domain of *Nk2* genes in the collar cord or the trunk nerve cords in *B. misakiensis.*

We additionally checked the distribution of serotonin-LIR neuronal components within the collar cord, because these neurons are restricted to the *Nkx2.1/2.2* domains in annelids and chordates. The serotonin-like immunoreactivity (LIR) nervous system of *B. misakiensis* has been described earlier [18], but the precise position of serotonin-LIR neurites within the collar cord has remained unknown. In the juvenile enteropneust serotonin-LIR neurons are present in the epidermis throughout all three body regions, with higher concentrations of somata in the proboscis and collar epidermis (Fig. 3O). Serotonin-LIR neurites form a basiepidermal nerve plexus in the proboscis and collar region. In the trunk region the serotonin-LIR neurites are condensed within the dorsal and ventral midline, in regions that constitute the nerve cords [18]. The neurulated collar cord passes through the mesocoel and is composed of a dorsal area of cells and a ventral neuropil (Fig. 3P, Q). The dorsal sheath of cells of the collar cord is devoid of serotonin-LIR somata (Fig. 3Q). Only two ventrolateral serotonin-LIR neurite bundles pass through the ventral neuropil. The lateral neurite bundles run adjacent to a pair of longitudinal muscle bundles that flank the collar cord (Fig. 3Q).

## Discussion

We investigated the expression domains of several genes involved in axial as well as mediolateral patterning of the nervous system of the indirect developing enteropneust *Balanoglossus misakiensis.* By using the pan-neuronal marker *Elav* for differentiating neurons [20–22], we found that the major parts of the central nervous system already develop in metamorphosing larvae prior to settlement. Within the collar region, the neural plate of the future collar cord is present and still part of the epidermis. In juveniles of *B. misakiensis*, the neural plate has neurulated completely to form the subepidermal tubular collar cord as also reported in other enteropneust species [10, 11, 16].

### Gene expression patterning of the collar cord in Enteropneusta

The transcription factors *Six3/6, Otx* and *engrailed* have been shown to play a conserved role in anteroposterior patterning and regionalization of the nervous system in chordates and in many other bilaterians [3]. It has been reported that *Six3/6* patterns the anteromost region of the nervous system in numerous animals [3, 23, 24]. We found that in *B. misakiensis* the expression pattern of *Six3/6* is likewise at the anteriormost region of the animal, while *Otx* and *engrailed* form circular epidermal domains around the collar and the posterior margin of the collar region, respectively (Fig. 4B’). These expression domains are spatially similar to what has been described in the spengelid *Schizocardium californicum* [15] as well as the harrimaniid enteropneust *Saccoglossus kowalevskii* (Fig. 4C’, taken from [13, 14]) Accordingly, we suggest a conserved role of neuronal as well as body region patterning for *Six3/6, Otx* and *engrailed* in Enteropneusta that is independent from their mode of development (direct vs. indirect, Fig.4B-C”). It is moreover a plesiomorphic feature for Enteropneusta that has been inherited from a common bilaterian ancestor [3, 13, 23].

**Fig. 4.**
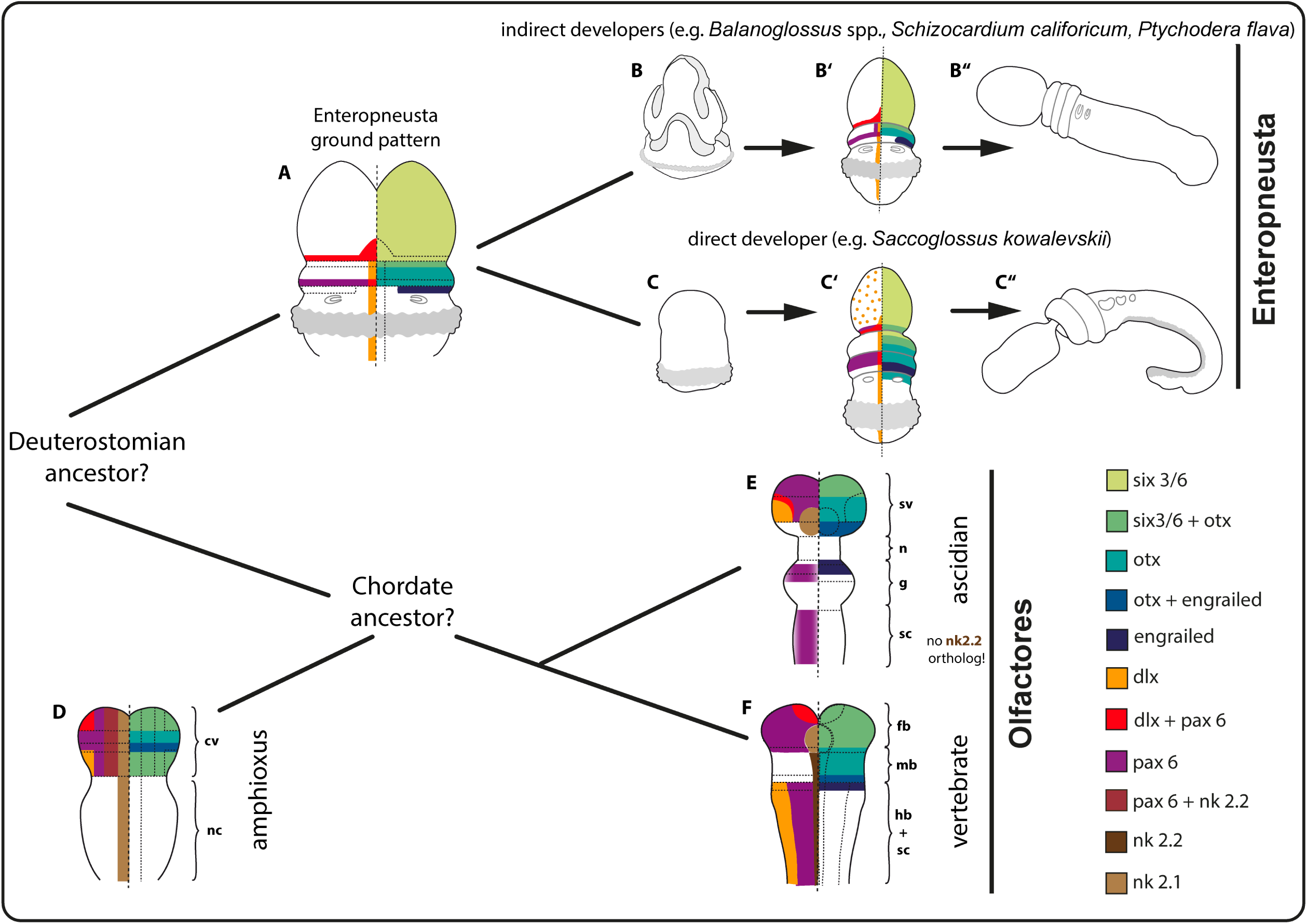
Comparison of axial patterning genes in the neural plate of diverse deuterostomian taxa with focus on different developing modes in enteropneusts. Neuronal patterning in enteropneusts is highly conserved and independent of the mode of development. The ancestral condition of mediolateral patterning for Deuterostomia remains elusive. See text for discussion. **A** Expression domains of the hypothetical enteropneust ancestor. **B-B’’** Selected developmental stages of *B. misakiensis*. **B** Metschnikoff larval stage. **B’** Metamorphosing Agassiz larval stage (this study). **B”** Juvenile worm. **C-C”** Selected developmental stages of *S. kowalevskii*. **C** Torpedo embryo stage. **C’** 1-gill slit hatchling (after data from [13, 14]). **C’’** Juvenile worm. **D** Expression domains in the neural plate of *Branchiostoma floridae* (after data from [33–35, 37, 38, 51]). **E** Expression domains in the neural plate of the ascidian *Ciona intestinalis* (after data from [33, 39, 52–54]). **F** Expression domains in the neural plate of the vertebrate *Mus musculus.* Scheme modified after [2] (data taken from [3, 14, 29, 36]). Note, all expression patterns are symmetrical, but are shown on one side only for clarity. cv = cerebral vesicle, fb = forebrain, g = ganglion, hb = hindbrain, mb = midbrain, n = neck, nc = nerve cord, sc = spinal cord, sv = sensory vesicle.

Next, we examined the expression pattern of *Dlx, Pax6* and *Nk2.1/2.2.* These transcription factors form mediolateral neurogenic domains in the neural tube of mouse, fruit fly as well as in the annelid *Platynereis dumerilii* [17, 25]. Our analysis in *B. misakiensis* shows that *BmiDlx* is expressed in a narrow longitudinal stripe in the dorsal midline of the neural plate in *B. misakiensis* (Fig. 4B’). A similar pattern has been reported for *Dlx* in *S. kowalevskii* (Fig. 4C’, after [14, 26]), *S. californicum* [15] and *Balanoglossus simodensis* [16], suggesting a conserved role of this transcription factor in neurogenesis in Enteropneusta. We then verified the expression pattern of *BmiPax6* and found that it forms two lateral stripes along the neural plate of *B. misakiensis* (Fig. 4B’). We show that *BmiPax6* is only expressed for a short period in the neural plate during metamorphosis and is entirely absent in early juveniles (2 d ps) (Fig. 3C, D). This is the only report of a distinct expression pattern of *Pax6* in the neural plate of an enteropneust species. In a comparable developmental stage of *S. kowalevskii* (1-gill-slit stage), *Pax6* is expressed in corresponding circular domains (Fig. 4C’, after [13, 14]), yet details from the neural plate are unknown. In *B. simodensis* and *S. californicum*, *Pax6* expression was not detected in the neural plate [15, 16]. Thus, *Pax6* expression in the collar cord might be a species-specific acquisition of *B. misakiensis* and not part of the enteropneust ground pattern (Fig. 4A).

Expression pattern analysis of the median progenitor markers *Nk2.1* and *Nk2.2* revealed that there is no expression domain of either of the *BmiNk2* genes in the developing neural plate or, later, in the collar cord in *B. misakiensis.* Instead, the main domains of *Nk2.1* and *Nk2.2* are detected in the pharyngeal endoderm (Fig. 3I-N). In the direct developer *S. kowalevskii* a likewise endodermal expression of both genes has been reported previously [13, 26] and in *Ptychodera flava Nk2.1* shows similar domains [27], suggesting a more general role in endoderm specification of these genes in enteropneusts [28]. Moreover, serotonergic neurons in vertebrates are usually restricted to the progenitor domains of *Nk2.1/2.2* [29]. Our data show that there is no median *Nk2.2* domain in the collar cord in *B. misakiensis.* Concordantly, no serotonin-LIR somata are present in the collar cord of *B. misakiensis.* In fact, *Nk2.2* does not co-localise with serotonin-LIR neurons in *B. misakiensis*. It was shown that serotonin-LIR neurons are indeed present in enteropneusts, yet all of them comprise bipolar neurons throughout the epidermis of *B. misakiensis, S. kowalevskii* [18] as well as *P. flava* [12]. Concluding so far, except for *Pax6,* the expression patterns of all the investigated genes in this study are highly congruent among the enteropneusts *S. kowalevskii, S. californicum, B. simodensis, P. flava* as well as *B. misakiensis,* which is why a similar function appears most likely. The data further reveals that neuronal patterning in the different families of Enteropneusta (Harrimaniidae, Spengelidae and Ptychoderidae) is not affected by different developmental modes. This conclusion is also supported by morphogenetic data of the developing nervous system in enteropneusts [18]. On that basis, we propose that a similar collar cord patterning was present in the last common ancestor of Enteropneusta (Fig. 4A). These results further corroborate the suitability of indirect as well as direct developing enteropneusts for serving as model organisms to conduct ‘evodevo’ studies in hemichordates.

### Comparative aspects of neural tube patterning among deuterostomes

Morphological similarities between the tubular collar cord and the chordate neural tube have not gone unnoticed and have been acknowledged from early on [30]. Therefore, we compare here the gene expression patterns of the studied transcription factors among different deuterostomes and discuss evolutionary implications.

Chordata comprises three major taxa, Cephalochordata, Tunicata and Vertebrata, of which the latter two form the monophyletic Olfactores [31, 32]. All three groups share corresponding expression domains of the transcription factors *Six3/6, Otx* and *Engrailed (En)* (Fig. 4D-F), which are restricted to the anterior portion of the neural plate, i.e., the future brain region [3, 33]. Thereby, coexpression of *Otx* and *En* mark the midbrain-hindbrain boundary (MHB) in vertebrates and the posterior margin of the sensory vesicle (brain) in the ascidian *Ciona intestinalis*. In contrast, the coexpressing domain of *Otx* and *En* in amphioxus is located in the midlevel of the brain region, whereas a second expression domain of *Six3/6* is present at the posterior end of the cerebral vesicle (Fig. 4D) [3, 34]. Moreover, all three groups show a median/ventral *Nk2.1* domain and expression domains of *Pax6* and *Dlx* in the brain region [13, 17, 29, 35–39]. Thus, the chordate ancestor likely had a similar brain patterned by these transcription factors [3].

Mediolateral patterning of the postcerebral part of the neural tube by *Pax6, Dlx* and *Nk2.1/2.2* differs considerably between chordates and needs further attention. The specific arrangement of lateral *Dlx*, mediolateral *Pax6* and median *Nk2* domains has been reported from the vertebrate spinal cord and hindbrain levels (posterior to MHB) as well as the annelid and insect ventral nerve cord (postcerebral) [2, 3]. The median column of *Nk2.2* is an exception as its domain projects anteriorly throughout the midbrain region and is replaced by *Nk2.1* in the vertebrate forebrain (Fig. 4F). However, ascidians share only a mediolateral *Pax6* domain with vertebrates, while *Dlx* and *Nk2.2* expression is absent from the postcerebral neural tube (Fig. 4E) [3, 39, 40]. Ascidians belong to Tunicata, a taxon of rapidly evolving animals with reduced genome size that have lost about 25 genes involved in developmental patterning including *Gbx, Wnt1* and *Nk2.2* [33, 41, 42]. Thus, the aberrant and lacking expression domains compared to vertebrates might be explained by secondary gene losses in Tunicata. Compared with this, amphioxus does not appear to be rapidly evolving. In fact, cephalochordates have retained all of the putative ancestral bilaterian homeobox genes [33, 42] and the genome of amphioxus is supposed to represent the most ancestral one among chordates, in parallel to a less derived morphology [42]. However, *Pax6* and *Dlx* expression are absent from the nerve cord in amphioxus, instead the median *Nk2.1* domain extends throughout the posterior neural plate (Fig. 4D) [38]. It should be mentioned here, that a median *Nk2.1/2.2* domain, a mediolateral *Pax6* as well as a lateral *Dlx* expression domain is very well present in amphioxus, yet these expression domains are located in the posterior region of the cerebral vesicle (Fig. 4D) and not in the postcerebral CNS as in vertebrates (Fig. 4D, F) and the protostomes *Platynereis dumerilii* and *Drosophila melanogaster* [2, 17]. Taken together, mediolateral expression domains of *Pax6*, *Dlx* and *Nk2* genes differ considerably among chordates, making it difficult to postulate a ground pattern for the last common chordate ancestor.

Since vertebrates and annelids (which exhibit a similar mediolateral patterning) are only distantly related taxa and comparable data from most intermediate groups are missing (see also [5]), an outgroup comparisons with Enteropneusta is most reasonable; not least, because this taxon is part of the Ambulacraria, the sister group of Chordata [32]. Comparison of the expression domains of *Six3/6, Otx* and *En* leads to the suggestion that the collar cord in enteropneusts might correspond to a region of the chordate brain rather than to the postcerebral neural tube (Fig. 4A, D-F). This is also supported by comparative Hox gene expression analysis in *S. kowalevskii* [3, 13]. In Enteropneusta the neural plate is patterned medially by *Dlx* ([13, 14, 16] this study) (Fig. 4A), whereas *Dlx* expression is restricted to the very lateral area of the brain in amphioxus and ascidians (Fig. 4D, E) and the spinal cord in vertebrates (Fig. 4F). Accordingly, there is no corresponding mediolateral patterning present in the enteropneust nervous system, and compared to chordates the expression domains of *Dlx* and *Pax6* are flipped in *B. misakiensis* (Fig. 4B’, D-F). These incongruent expression patterns might be explained by the fact that dorsoventral signaling of *Bmp* and *chordin,* which is responsible for the placement of the mediolateral patterning domains, is inverted in enteropneusts and chordates [26]. It was shown in *S. kowalevskii* that the tubular collar cord develops from the *Bmp*-expressing side, whereas the dorsal neural tube of chordates and the ventral nerve cord of protostomes form at the Chordin-expressing side [26, 43]. Concordantly, markers of midline cells in the chordate neural tube such as *Sim* and *Netrin* are expressed in the ventral ectoderm in enteropneusts while lateral markers of the chordate neural tube such as *Dlx* are expressed in the dorsal ectoderm ([26], this study). Thus, according to the dorsoventral (D-V) inversion hypothesis [44, 45], the collar cord might in fact be positioned on the “wrong side”, that is, corresponding to the ventral side of chordates.

## Conclusion

A complex mediolateral patterning of the postcerebral nervous system by *Pax6*, *Dlx* and *Nk2.1/Nk2.2,* among others, has been reported from vertebrates and the protostomes *Platynereis dumerilii* and *Drosophila melanogaster* [2, 3, 17]. However, comparison of their expression domains among different deuterostome taxa does not suggest that a likewise patterned postcerebral nervous system was present in the last deuterostomian ancestor (Fig. 4). Moreover, the tubular collar cord of Enteropneusta shows no expression domains of *Dlx* or *Pax6* that clearly correspond to the chordate brain. The “flipped” domains of *Dlx* and *Pax6* in enteropneusts are likely the result of an inverted *BMP/Chordin* expression compared to chordates [26], and suggest that the collar cord represents an independent acquisition of Enteropneusta, thereby contrasting the results from ultrastructural and classical investigations [10, 11]. Accordingly, the question of homology versus independent evolution of the enteropneust collar cord and the chordate neural tube might primarily depend on the (subjective) decision whether one favors the morphogenetic over the molecular (gene expression) evidence or vice versa. While this may sound frustrating at first, the incongruence in the dataset currently available should instead motivate today’s developmental biologists to further engage in research into this area in order to finally settle one of the key issues in animal evolution: the origin of bilaterian centralized nervous systems.

## Materials & Methods

### *Balanoglossus misakiensis* (Kuwano, 1902)

Adult *B. misakiensis* were collected at a depth of 1 to 2 m at Sunset beach, Aomori-Bay, Asamushi, Aomori, Japan, in June 2012 and June 2014. Specimens were transported to the Research Center for Marine Biology Tohoku University in Asamushi and were kept in aquaria with running filtered seawater at ambient water temperature (24 - 26°C) as previously described [18, 46]. Spawning, in vitro fertilization, and fixations were performed as described earlier [18].

### Immunolabelling and confocal laser scanning microscopy

Juveniles of *B. misakiensis* (2-gill-slit juvenile = 3 days post settlement) were fixed with 4% paraformaldehyde (PFA) in phosphate buffer (PBS). Specimens were processed using standard protocols as previously described [18].

### RNA extraction, transcriptome analysis and gene cloning

More than 1,000 larvae from developmental stages of *B. misakiensis* ranging from early hatched tornaria to three day old juvenile worms were fixed in RNAlater (Sigma). Total RNA was extracted from a mix of developmental stages using RNeasy Mini Kit from Qiagen. Extracted RNA was sent to Eurofins (Germany) for Illumina HiSeq 2000 sequencing using paired-end read module resulting in reads of 100bp length. Obtained reads were assembled to contigs using Trinity software under standard parameters and the transcriptome was analysed for sequences of interest with BLAST search in Geneious 6.1 (Biomatters, New Zealand). Primers were generated to obtain fragments of *Elav, Six3/6, Pax6, Dlx, Otx, Engrailed, Nk2.1* and *Nk2.2* (for primer sequences and accession numbers see supplemental material) in order to sub-clone into pGemT Easy vector (Promega).

### Phylogenetic analysis

Full protein sequences were aligned using MUSCLE and Regions with low-quality alignments for the Elav phylogenetic anaylsis were trimmed by TrimAl 1.2 rev 59 [47]. ProtTest 2.4 [48] analysis retrieved LG (+G+F) and JTT (+I+G+F) for the Elav (Fig. S2) and the homeobox protein (Fig. S3) analyses respectively as best-fitting models for the phylogenetic reconstruction. The maximum likelihood tree was then generated with PhyML 3.0 ([49], BIONJ input tree, optimised tree topology, 4 substitution rate categories, best of NNI and SPR, 100 non-parametric bootstrap replicates).

### Probe synthesis and *in situ* hybridization

Chromogenic *in situ* hybridizations were performed on whole-mounts following the protocol from Röttinger and Martindale [50] with minor adjustments for *B. misakiensis.* Metamorphosing larvae (Agassiz stage) and juveniles (2-gill-slit stage) were treated with 10 ng/μl Proteinase K (Roth) for 4 min at room temperature. Colour development was stopped by three washes in PTw (phosphate buffered saline + 0.1% Tween20) and postfixed with 4% PFA for 1 hour. Animals were transferred into 100% EtOH over night for clearing and mounted in 80% glycerol.

### Controls

Controls with sense probes were conducted in order to identify unspecific binding and probe trapping during *in situ* hybridization. The protocoel within the proboscis region turned out to be a perfect trap for any probe (Fig. S1A). Perforation of the proboscis using a thin tungsten needle helped to solve this problem (Fig.S1D, E). However, probe trapping could not always be eliminated, which is why a blue protocoel persists in the in situ hybridizations of *Dlx* (Fig.3E-H) and in *Otx* and *engrailed* in juveniles (Fig.2G, H, K, L).

## Acknowledgements

We thank Takuya Minokawa from the Research Center for Marine Biology Tohoku University in Asamushi, Japan for, providing laboratory space and equipment. This study was funded by a Lise-Meitner grant from the Austrian Science Fond (FWF) to SK-S (M 1485-B19). The collection trip of SK-S to Japan was financially supported by a stipend from the Research Center for Marine Biology, Tohoku University, Japan. Tim Wollesen (University Vienna) is thanked for his help in basic molecular biological methods. SK-S thanks Thomas Eder and Thomas Rattei (both University of Vienna) for their kind assistance with Illumina transcriptome assembly. SK-S kindly acknowledges the valuable support of Eric Röttinger (University of Nice) and Grigory Genikhovich (University of Vienna) that allowed the establishment of a reliable in situ protocol for *B. misakiensis*. Patrick Steinmetz (Sars Center, Bergen) is thanked for critical comments and suggestions on earlier versions of this manuscript.

## Author Contributions

SK-S and AW designed the study. MU and SK-S collected and cultured material. SK-S conducted IHC and cLSM analyses. SK-S extracted RNA, assembled the transcriptome, cloned all gene sequences, and performed in situ hybridizations. DP aligned sequences, conducted phylogenetic analyses and built orthology trees of the genes. SK-S wrote the manuscript with input from AW. All authors read, provided input, and approved the final version of the manuscript.

## Accession codes

All gene sequences will be deposited at GenBank upon acceptance of this manuscript.

## Competing interests

The authors declare that they have no competing financial interests.

## References

1. Holland ND (2003) Early central nervous system evolution: An era of skin brains? Nat Rev Neurosci 4:1–11.

2. Arendt D, Denes AS, Jekely G, Tessmar-Raible K (2008) The evolution of nervous system centralization. Phil Trans R Soc B 363: 1523–1528.

3. Holland LZ, Carvalho JE, Escriva H, Laudet V, Schubert M, Shimeld S, Yu J-K (2013) Evolution of bilaterian central nervous systems: a single origin? EvoDevo 4:27.

4. Arendt D, Tosches MA, Marlow H (2015) From nerve net to nerve ring, nerve cord and brain – evolution of the nervous system. Nature Rev Neurosci 17: 61–72.

5. Hejnol A, Lowe CJ (2015) Embracing the comparative approach: how robust phylogenies and broader developmental sampling impacts the understanding of nervous system evolution. Phil. Trans. R. Soc. B 370: 20150045.

6. Lowe CJ, Clarke DN, Medeiros DM, Rokhsar DS, Gerhart J (2015) The deuterostome context of chordate origins. Nature 520: 456–465.

7. Kaul-Strehlow S, Röttinger E (2015) Hemichordata in: Evolutionary developmental biology of invertebrates Vol. 6 (ed. Andreas Wanninger). Springer Verlag, Berlin.

8. Bullock TH (1946) The anatomical organization of the nervous system of Enteropneusta. Q J Microsc Sci 86: 55–112.

9. Knight-Jones EW (1952) On the nervous system of *Saccoglossus cambrensis* (Enteropneusta). Philos Trans R Soc Lond B 236: 315–354.

10. Morgan TH (1894) The development of Balanoglossus. J Morphol 9: 1–86.

11. Kaul S, Stach T (2010) Ontogeny of the collar cord: Neurulation in the hemichordate *Saccoglossus kowalevskii.* J Morph 271: 1240–1259.

12. Nomaksteinsky M, Roettinger E, Dufour HD, Chettouh Z, Lowe CJ, Martindale MQ, Brunet J-F (2009) Centralization of the Deuterostome Nervous System Predates Chordates. Curr Biol 19:1264–1269.

13. Lowe CJ, Wu M, Salic A, Evans L, Lander E, Stange-Thomann N, Gruber CE, Gerhart J, Kirschner M. (2003) Anteroposterior patterning in hemichordates and the origin of the chordate nervous system. Cell 113: 853–865.

14. Pani A, Mullarkey EE, Aronowicz J, Assimacopoulos S, Grove EA, Lowe CJ (2012) Ancient deuterostome origins of vertebrate brain signalling centres. Nature 438: 289–295.

15. Gonzalez P, Uhlinger KR, Lowe CJ (2017) The Adult Body Plan of Indirect Developing Hemichordates Develops by Adding a Hox-Patterned Trunk to an Anterior Larval Territory. Curr Biol 27: 87–95.

16. Miyamoto N, Wada H (2013) Hemichordate neurulation and the origin of the neural tube. Nat Commun 4:2713.

17. Denes AS, Jékely G, Steinmetz PRH, Raible F, Snyman H, Prud’homme B, Ferrier DEK, Balavoine G, Arendt D (2007) Molecular architecture of annelid nerve cord supports common origin of nervous system centralization in Bilateria. Cell, 129:277–288.

18. Kaul-Strehlow S, Urata M, Minokawa T, Stach T, Wanninger A (2015) Neurogenesis in directly and indirectly developing enteropneusts: of nets and cords. Org. Divers. Evol. 15, 405–422. (doi:10.1007/s13127-015-0201-2)

19. Nielsen C, Hay-Schmidt A (2007) Development of the enteropneust *Ptychodera flava:* ciliary bands and nervous system. J Morphol 268:551–570.

20. Soller, M., and White, K. (2004). Elav. Curr. Biol. 14, R53.

21. Berger, C., Renner, S., Lüer, K., Technau, G.M., 2007. The commonly used marker ELAV is transiently expressed in neuroblasts and glial cells in the Drosophila embryonic CNS. Dev. Dyn. 236, 3562–3568.

22. Pascale, A., Amadio, M., Quattrone, A. (2008). Defining a neuron: neuronal ELAV proteins. Cell Mol Life Sci 65: 128–140.

23. Steinmetz, P. R. et al. Six3 demarcates the anteriormost developing brain region in bilaterian animals. Evodevo 1, 14 (2010).

24. Marlow H et al.: Larval body patterning and apical organs are conserved in animal evolution. BMC Biology 2014 12:7.

25. Holland LZ (2015) The origin and evolution of chordate nervous systems. Phil. Trans. R. Soc. B 370: 20150048. http://dx.doi.org/10.1098/rstb.2015.0048

26. Lowe CJ, Terasaki M, Wu M, Freeman RM, Runft L, Kwan K, Haigo S, Aronowicz J, Lander E, Gruber C, Smith M, Kirschner M, Gerhart J (2006) Dorsoventral patterning in hemichordates: insights into early chordate evolution. PLoS Biol 4: e291.

27. Takacs, CN., Moy, VN, Peterson, KJ (2002) Testing putative hemichordate homologues of the chordate dorsal nervous system and endostyle: expression of *NK2.1* (TTF-1) in the acorn worm *Ptychodera flava* (Hemichordata, Ptychoderidae). Evol &Dev 4: 405–417.

28. Okkema, P.G., Ha, E., Haun, C., Chen, W., and Fire, A. (1997) The *Caenorhabditis elegans* NK-2 homeobox gene ceh-22 activates pharyngeal muscle gene expression in combination with pha-1 and is required for normal pharyngeal development. Development 124, 3965–3973.

29. Shimamura, K., Hartigan, D. J., Martinez, S., Puelles, L. & Rubenstein, J. L. (1995) Longitudinal organization of the anterior neural plate and neural tube. Development 121: 3923–3933.

30. Bateson W (1884) The early stages of the development of Balanoglossus (sp. incert.). Q J Microsc Sci, NS 24: 208–236, pls 18-21.

31. Delsuc F, Brinkmann H, Chourrout D, Philippe, H (2006) Tunicates and not cephalochordates are the closest living relatives of vertebrates. Nature 439: 965–968.

32. Cannon JT, Vellutini BC, Ill JS, Ronquist F, Jondelius U, Jejnol A (2016) Xenacoelomorpha is the sister group to Nephrozoa. Nature 530: 89–93.

33. Castro LFC, Rasmussen SLK, Holland PWH, Holland ND, Holland LZ (2006) A Gbx homeobox gene in amphioxus: Insights into ancestry of the ANTP class evolution of the midbrain/hindbrain boundary. Dev Biol 295: 40–51.

34. Kozmik Z, Holland ND, Kreslova J, Oliveri D, Schubert M, Jonasova K, Holland LZ, Pestarino M, Benes V, Candiani S (2007) *Pax-Six-Eya-Dach* network during amphioxus development: Conservation *in vitro* but context specificity *in vivo.* Dev Biol 306: 143–159.

35. Holland ND, Panganiban G, Henyey EL, Holland LZ (1996) Sequence and developmental expression of *AmphiDll*, an amphioxus *Distal-less* gene transcribed in the ectoderm, epidermis and nervous system: insights into evolution of craniate forebrain and neural crest. Development 122: 2911–2920.

36. Mastick, G. S., Davis, N. M., Andrew, G. L. & Easter, S. S Jr. (1997) Pax-6 functions in boundary formation and axon guidance in the embryonic mouse forebrain. Development 124: 1985–1997.

37. Glardon S, Holland LZ, Gehring WJ, Holland ND (1998) Isolation and developmental expression of the amphioxus *Pax-6* gene (*AmphiPax-6*): insights into eye and photoreceptor evolution. Development 125: 2701–2710.

38. Venkatesh TV, Holland ND, Holland LZ, Su M-T, Bodmer R (1999) Sequence and developmental expression of amphioxus *AmphiNk2-1:* insights into the evolutionary origin of the vertebrate thyroid gland and forebrain. Dev Genes Evol 209: 254–259.

39. Irvine SQ, Cangiano MC, Millette BJ, Gutter ES (2007) Non-overlapping Expression Patterns of the Clustered *Dll-A/B* Genes in the Ascidian *Ciona intestinalis.* J Exp Zool B 308: 428–441.

40. Edvardsen RB, Seo H-C, Jensen MF, Mialon A, Mikhaleva J, Bjordal M, Cartry J, Reinhardt R, Weissenbach J, Wincker P, Chourrout D (2005) Remodelling of the homeobox gene complement in the tunicate *Oikopleura dioica*. Curr Biol 15: R12–R13.

41. Holland LZ (2009) Chordate roots of the vertebrate nervous system: expanding the molecular toolkit. Nat Rev 10: 736–746.

42. Holland ND (2010) From genomes to morphology: a view from amphioxus. Acta Zool 91: 81–86.

43. De Robertis EM (2008) Evo-Devo: Variations on Ancestral Themes. Cell 132: 185–195.

44. Geoffrey St-Hilaire E (1822) Considérations générales sur la vertèbre. Mémoires, Mus d’His Nat 9:89–119.

45. Arendt, D., and Nübler-Jung, K. (1999) Comparison of early nerve cord development in insects and vertebrates. Development 126, 2309–2325.

46. Urata M, Yamaguchi M (2004) The Development of the Enteropneust Hemichordate *Balanoglossus misakiensis* KUWANO. Zoological Science 21: 533–540.

47. Capella-Gutiérrez S, Silla-Martínez JM, Gabaldón T (2009) trimAl: a tool for automated alignment trimming in large-scale phylogenetic analyses. Phylogenetics 25: 1972–1973.

48. Abascal F, Zardoya R, Posada D (2005) ProtTest: selection of best-fit models of protein evolution. Phylogenetics 21: 2104–2105.

49. Guindon S, Dufayard J-F, Lefort V, Anisimova M, Hordijk W, Gascuel O (2010) New Algorithms and Methods to Estimate Maximum-Liklihood Phylogenies: Assessing the Performance of PhyML 3.0. Syst Biol 59: 307–321.

50. Röttinger, E. and Martindale, M. Q. (2011). Ventralization of an indirect developing hemichordate by NiCl suggests a conserved mechanism of dorso-ventral (D/V) patterning in Ambulacraria (hemichordates and echinoderms). Dev. Biol. 354: 173–190.

51. Holland LZ, Venkatesh TV, Gorlin A, Bodmer R, Holland ND (1998) Characterization and developmental expression of AmphiNk2-2, an NK2 class homeobox gene from amphioxus (Phylum Chordata; Subphylum Cephalochordata). Dev Genes Evol 208: 100–105.

52. Moret F, Christiaen L, Deyts C, Blin M, Joly J-S, Vernier P (2005) The dopamine-synthesizing cells in the swimming larva of the tunicate *Ciona intestinalis* are located only in the hypothalamus-related domain of the sensory vesicle. Europ J Neurosci 21: 3043–3055.

53. Hudson C, Lemaire P (2001) Induction of anterior neural fates in the ascidian *Ciona intestinalis.* Mech Dev 100: 189–203.

54. Mazet F, Hutt JA, Millard J, Shimeld SM (2003) Pax gene expression in the developing central nervous system of *Ciona intestinalis.* Gene Exp Patt 3: 743–745.

